# Maternal separation affects fronto-cortical activity in rat pups during dam-pup interactions and behavioral transitions

**DOI:** 10.1101/2021.05.19.444831

**Authors:** Manon Ranger, Tatyana B. Behring, Jasmine H. Kaidbey, Muhammad Anwar, Allie B. Lipshutz, Isabelle Mollicone, Gehad Hassan, Kathryn Fasano, Nicole K. Hinz, Robert J. Ludwig, Michael M. Myers, Martha G. Welch, Dani Dumitriu

**Affiliations:** Division of Developmental Neuroscience, Department of Psychiatry, Columbia University Irving Medical Center, New York, NY, 10032, USA; Nurture Science Program, Department of Pediatrics, Columbia University Irving Medical Center, New York, NY, 10032, USA; School of Nursing, University of British Columbia, Vancouver, BC, V6T 2B5, Canada; Sackler Institute for Developmental Psychobiology, Columbia University, New York, NY, 10032, USA; Division of Child and Adolescent Health, Department of Pediatrics, Columbia University Irving Medical Center, New York, New York, NY, 10032, USA

**Keywords:** Early-life stress, electrocorticography, cortical development, maternal separation, rat pups, maternal behavior

## Abstract

Early-life stress is known to impair neurodevelopment. Prior work from our group showed that prolonged physical and emotional separation necessitated by the medical needs of preterm infants (born <37 weeks) is associated with lower electroencephalogram (EEG) power in frontal areas, and that trend can be reversed by an intervention that enhances the physical and emotional contact between preterm infant and mother. Here we sought to model the changes in preterm infant frontal EEG power in a rodent model. We examined effects of daily maternal separation (MS) on frontal cortex electrophysiological (electrocorticography [EcoG]) activity in neonatal rats. We also explored the effects of dam-pup behavioral interactions on EcoG activity. From postnatal days (P) 2-10, rat pups were separated daily from their dam and isolated from their littermates for 3 hours. Control rats were normally reared. On P10, pups were implanted with telemetry devices and an electrode placed on the left frontal dura. EcoG activity was recorded during daily sessions over the next four days while pups remained in the home cage, as well as in response to a pup-dam isolation-reunion paradigm at P12. EcoG power was computed in 1 Hz frequency bins between 1-100 Hz. Dam and pup behavioral interactions during recording sessions were coded and synchronized to EcoG data. MS pups showed lower EcoG power during dam-pup interactions. These data parallel human and provide evidence of lower fronto-cortical activity as an early marker of early-life stress and possible mechanism for long-term effects of maternal separation on neurobehavioral development.

## Introduction

Early-life stress impairs neurodevelopment in both humans and animals (Lippmann et al., 2007; Lupien et al., 2009; Callaghan et al., 2014; Curley and Champagne, 2016; Tractenberg et al., 2016; Walker et al., 2017; Yang et al., 2017; Bonapersona et al., 2018). Maternal separation (MS) is one such stressor, resulting in profound impact on child physical and emotional health (MacLean, 2003; Nelson et al., 2013; Humphreys et al., 2015; Welch and Myers, 2016). In some cases, MS is unavoidable, such as when necessitated by the medical needs of infants born prematurely or with other medical needs necessitating admission to the neonatal intensive care unit (NICU). While other mechanisms, including prematurity itself, procedural pain, and hypoxic injury, contribute to deleterious long-term health outcomes in these infants, MS is thought to play a key role (Liu et al., 2016; Welch and Myers, 2016). An estimated 15 million babies are born prematurely every year globally (Chawanpaiboon et al., 2019) most of which require NICU admission. Since the MS associated with NICU hospitalization is one of few modifiable risk factors for the long-term outcomes of these infants it is imperative to both understand the mechanisms by which MS impact brain development and develop methods to mitigate this risk.

To this end, our translational research program at the Nurture Science Program (www.nurturescienceprogram.org) developed Family Nurture Intervention (FNI) (Welch et al., 2012), an intervention aimed at strengthening the emotional connection between mother and child. In a first randomized clinical trial (RCT) of FNI on premature infants admitted to the NICU, we showed neurodevelopmental benefits up to age 5 (Welch et al., 2015, 2020). Infants in the FNI group had significantly higher frontal EEG power than infants in the standard care group (Welch et al., 2014; Myers et al., 2015), pointing to altered brain maturation as a potential mechanism mediating the benefits of mother-infant early interactions. However, to fully investigate the mechanisms linking MS, maternal nurture, and brain development, an animal model is crucial. Our goal was therefore to develop a rodent MS model that recapitulates frontal EEG power differences observed in NICU infants with and without FNI. We hypothesized that MS rat pups would show lower frontal brain activity similar to NICU infants in the standard care condition, who experience relatively higher levels of MS than FNI infants.

Rodent MS is a well-established model of early-life stress and has been shown to induce multi-system alterations (Callaghan et al., 2014; Lippmann et al., 2007; Mooney-Leber and Brummelte, 2017; Tractenberg et al., 2016),including changes in prefrontal cortex myelination at weaning (Yang et al., 2017) and impaired prefrontal-hippocampal coupling (Reincke and Hanganu-Opatz, 2017). Prior work describing electrical activity in the developing rodent brain has primarily focused on cortical and subcortical local field potentials (LFP) (Sarro et al., 2014; Opendak et al., 2020). Here, given our explicit goal to model human infant EEG findings, we chosee lectrocorticography (EcoG) recordings on the dura over the left frontal cortex of rat pups, the brain area where we observed the largest differences in FNI versus standard care infants (Welch et al., 2014). Frontal EcoG power was measured in MS versus control pups at postnatal days (P) 11-14 after a standard 3-hour daily MS protocol from P2 to P10. We evaluated EcoG during (1) naturally occurring behaviors such as arched back nursing and licking and grooming, (2) during behavioral transitions such as milk ejections and dam coming into nest, and (3) during an isolation-reunion paradigm we previously tested to induce pup ultrasonic potentiation following the same MS protocol (Kaidbey et al., 2019).

## Methods

### Subjects and Maternal Separation Procedure

All procedures used in this study were reviewed and approved by the Columbia University and New York State Psychiatric Institute’s Institutional Animal Care and Use Committee. Nineteen timed-pregnant Sprague-Dawley female rats (obtained from Charles-River) arrived at our animal facility at E17-18 gestation where they were housed individually until delivery of pups on E22. Food and water were provided ad libitum and rooms were maintained on a 12:12 hr darklight cycle (lights on at 7 am).

Our separation paradigm was based on well-established models (reviewed in (Gutman and Nemeroff, 2002; Pryce and Feldon, 2003; Molet et al., 2014; Tractenberg et al., 2016)) and our prior work showing this MS protocol induces alterations in USV potentiation during an isolation-reunion protocol (Kaidbey et al., 2019). On P2, pups were weighed, litters culled to 8 pups (1:1 male-to-female ratio when possible), and randomly assigned to MS versus normally reared controls. Pups in the MS group were subjected to daily 3-hour separation from P2 thru P10 as follows: each day at 10am dams were removed from the home cage and placed into a novel cage in a separate room with access to food and water; pups were weighed and placed individually in small plastic cylinders (Figure 1A). The eight cylinders were placed in a clean cage which was then put in an incubator kept at constant temperature (P2-4 at 34°C, P5-7 at 33°C, P8-10 at 32°C) (based on (Satinoff, 1991; Plotsky et al., 2005; Callaghan and Richardson, 2011; Chocyk et al., 2013)). At 1pm, pups were returned to their home cage and reunited with their dam. The home cage was returned to the animal housing room and left undisturbed until the following day. Control litters were left undisturbed in their home cages with their respective dams, except for daily weighing.

**Fig. 1.**
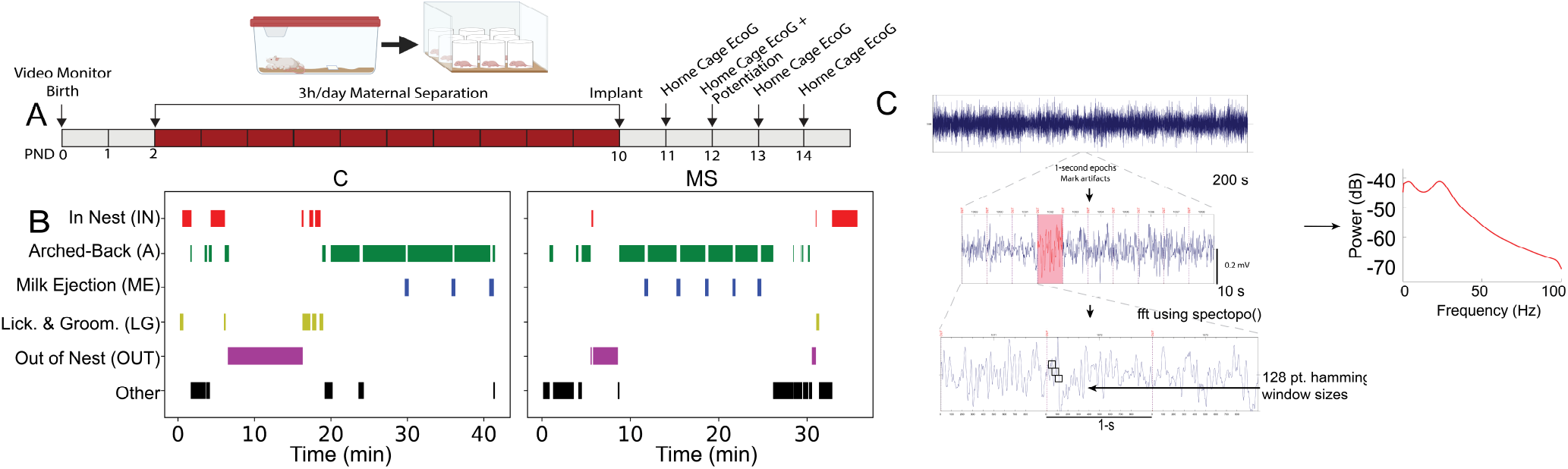
The maternal separation paradigm and the home cage behavioral assessment used for EcoG preprocessing and analysis. **(A)** Study timeline of the MS paradigm, surgery, and home cage testing. Pups were video monitored for birth (P0) and 3h/day MS lasted from P2 to P10. On P10 EEG telemetry devices were implanted. Home cage EcoG and behavioral data were collected from P11 to P14. A maternal potentiation paradigm took place on P12. **(B)** Raster plots of representative C (left) and MS (right) behavioral coding traces with each behavioral epoch plotted over time. **(C)** Behavioral epoch durations expressed as a percentage of total recording time (% Time) from P11 to P14. P11: C (*n* = 8), MS (*n* = 16); P12: C (*n* = 11), MS (*n* = 17); P13: C (*n* = 9), MS (*n* = 16); P14: C (*n* = 9), MS (*n* = 15). **(D)** EcoG preprocessing steps: 200-sec EcoG snippet of a trace that would undergo 1-sec epoching and artifact rejection. Fast Fourier transform (FFT) (spectopo()) on each epoch was used to generate power spectral density data. EcoG, electrocorticography; MS, maternal separation group; C, Control group; P, postnatal day.

### EcoG Surgical Implantation

On P10, two pups (1 male/1 female) from each litter were surgically implanted with a 1.6 g wireless single channel biopotential telemetric device (PhysioTel® ETA-F10, Data Science International) connected to an EEG electrode placed directly on the dura over the left frontal cortex allowing recordings of electrocorticography (EcoG). Pups were anesthetized with isoflurane and the surgical procedure was performed using aseptic technique. Anesthetized animals were kept warm on a water pad until they had regained righting reflex post-surgery. A 1 cm incision was made in the scalp along the dorsal midline from the posterior margin of the eyes to a point midway between the scapulae. The scalp was exposed and skull dried. One hole was drilled for the recording electrode using coordinates to target the left frontal cortex (from Bregma: 2.4 mm anterior and 1.5 mm medio-lateral) and a second hole for the reference electrode over the right hemisphere. A Teflon-coated 0.18 mm diameter stainless steel electrode was inserted in each hole, placed directly over the dura, and secured in place with dental cement. A subcutaneous pocket was made dorsally to allow placement of the telemetric transmitter. The skin incision was closed with 10-0 prolene non-absorbable sutures (Ethicon) and anesthesia was discontinued but supplemental warmth (water pad) was continued until the pup was fully awake. Post-surgical analgesia with carprofen was administered immediately after surgery and continued daily for 3 days. When fully recovered from anesthesia and mobile, pups were returned to their home cage; they were monitored throughout the post-operative phase for any signs of distress, pain, or infection. Implant surgeries were less than 45 min. Pups *n* = 11 C [4:7, male:female]; *n* = 17 MS [9:8, male:female]) included in the study recovered well, without signs of trauma or infection, moving freely and feeding normally within 10-15 min post-operatively, and appearing normal throughout the course of the experiment until sacrificed at P14-16 for surgical extraction of telemetry device. Pups that showed signs of poor recovery with minimal or no daily weight gain, infections, or that died anytime during the experiment (*n* = 22 C [7:4, male:female] and *n* = 8 MS [5:3, male:female]) were excluded from all analyses even if some recordings had been obtained.

### EcoG Data Collection

From P11 to P14, EcoG activity was recorded for 15-50 minutes daily between 9 am and 1 pm in freely moving pups, during normal interactions with their dam and littermates within their home cage. Before each recording session, the litter and dam were left alone and undisturbed in the room for a minimum of 30 min to allow them to acclimatize to the recording environment and thereby promote normal behavior. EcoG data were acquired using LabChart software (ADInstruments) with synchronized video recording via a webcam for subsequent behavioral coding of dam-pup interactions. The EEGLAB MATLAB toolbox and Signal Processing Toolbox (Delorme and Makeig, 2004) were used for preprocessing and analysis of the EcoG recordings (see EcoG data processing section).

### Dam/pup Behavior Coding During EcoG Recordings

During daily EcoG recording sessions, litters from both groups were observed to quantify undisturbed dam-pup home cage behavioral interactions. We used previously validated methods and established checklists (Myers et al., 1989; Franks et al., 2011; Pena et al., 2014) to categorize and measure the duration of various maternal behaviors during the 15-50 min continuous video recordings. Video recordings captured during each session were coded by 4 observers who were blinded to the treatment groups. The durations of the following five behaviors were logged: dam in-nest (IN) – the dam is in the nest with the pups; dam out-of-nest (OUT) – dam is not in contact or present with her pups within the nest but remains in the home cage; Arched Back Nursing (A), which included : i) high-arched nursing - dam is in a high-arched posture over pups with all four legs splayed and ii) low-arched nursing - dam is over the litter, but is not fully arched and with no extension of her legs; licking and grooming (LG) - dam is licking the pup (either anogenital or whole body); milk ejection (ME) – nipple attached pups display a wave of dramatic stretching movements and other motor activity (e.g. switching to a neighboring nipple) before going back to their original quiet state. Note that licking-grooming behaviors could co-occur with low arched-back nursing. The exact start/stop time of each specific behavior of interest was coded using BORIS (Friard et al., 2016). Inter-rater reliability for behavior coding was maintained above 0.9 (Cohen’s Kappa) throughout the study.

### Maternal Potentiation of Behavioral Reactions to Isolation Test

On P12, all pups, including those that had the EEG implant, went through behavioral testing for maternal potentiation of pup ultrasonic vocalization (USV). This procedure was based on a well-established paradigm consisting of pup isolation, brief reunion with dam, and a second isolation during which rates of USV production are increased over the rates during the initial isolation (Hofer et al., 2002). Methods and USV related findings have been described elsewhere (Kaidbey et al., 2019). Briefly, between 10 am and 12 pm, 30-minutes before testing, the dam was removed from the home cage and brought to a novel room in a cage with fresh bedding, water and food. The home cage with the pups was placed on a heating pad in a separate novel room. Upon completion of the 30-minute separation, each pup, in succession and random order, was gently brought to the testing room (22.3°*C*) in a novel small (25 × 14 × 11.5 cm) clear polycarbonate rectangular box (no bedding). Each pup was observed in the novel container during three conditions: first isolation of 3 min (ISO1), a 3 min reunion with their dam (REUN), and finally a second 3 min isolation (ISO2). USVs were recorded with an ultrasound sensitive microphone connected to a computer via UltraSoundGate 116 USB audio device (Avisoft Bioacoustics, Glienicke, Germany) placed 2 inches above the test cage, but are not reported here. Upon completion of the second isolation, each pup was returned to their home cage and reunited with their littermates. Dams were returned to their home cages and reunited with their litter once all pups had gone through the potentiation paradigm. EcoG and behavior were recorded throughout the three phases of the potentiation paradigm in the surgically implanted pups. Dam-pup behavior during REUN was coded as either being In-Contact or Not-in-Contact and EcoG power was only examined for the periods when the pup was In-Contact with the dam.

### Maternal Potentiation of Behavioral Reactions to Isolation Test

On P12, all pups, including those that had the EEG implant, went through behavioral testing for maternal potentiation of pup ultrasonic vocalization (USV). This procedure was based on a well-established paradigm consisting of pup isolation, brief reunion with dam, and a second isolation during which rates of USVs production are increased over the rates during the initial isolation (Hofer et al., 2002). Methods and USV related findings have been described elsewhere (Kaidbey et al., 2019). Briefly, between 10am and 12pm, 30-minutes before testing, the dam was removed from the home cage and brought to a novel room in a cage with fresh bedding, water and food. The home cage with the pups was placed on a heating pad in a separate novel room. Upon completion of the 30-minute separation, each pup, in succession and random order, was gently brought to the testing room (22.3°C) in a novel small (25 × 14 × 11.5 cm) clear polycarbonate rectangular box (no bedding). Each pup was observed in the novel container during three conditions: first i solation o f 3 min (ISO1), a 3 min reunion with their dam (REUN), and finally a second 3 min isolation (ISO2). USVs were recorded with an ultrasound sensitive microphone connected to a computer via UltraSoundGate 116 USB audio device (Avisoft Bioacoustics, Glienicke, Germany) placed 2 inches above the test cage, but are not reported here. Upon completion of the second isolation, each pup was returned to their home cage and reunited with their littermates. Dams were returned to their home cages and reunited with their litter once all pups had gone through the potentiation paradigm. EcoG and behavior were recorded throughout the three phases of the potentiation paradigm in the surgically implanted pups. Dam-pup behavior during REUN was coded as either being In-Contact or Not-in-Contact and EcoG power was only examined for the periods when the pup was In-Contact with the dam.

### EcoG data processing

EcoG recordings were band-pass filtered from 0.1 to 100 Hz and down sampled from 1000 to 512 samples for Fast Fourier Transforms (FFTs). For each behavior, one-second EcoG epochs were created, artifacts were marked and then automatically rejected using the EEGLAB rejepoch function. FFT power analysis using the spectopo function was performed on the filtered and artifact free EcoG data for each 1-sec epoch of isolated behavioral state. Oscillatory power was computed for 100 frequency bands between 1 and 100 Hz (1 Hz bins). We quantified EcoG power during Out of Nest (OUT), In-Nest (IN), Arched Back Nursing (A), Licking and Grooming (LG), and Milk Ejection (ME) behavioral states. The same EcoG data processing was also applied to quantify EcoG power within each phase of the maternal potentiation testing paradigm. In the reunion phase, we only included EcoG data when pups were In-Contact with their dams. Sample size varied between P11, P12, P13 and P14, thus, EcoG data was analyzed separately for each day.

### Statistical Analysis

For each recording day (P11 through P14) and session (home cage and potentiation), repeated measures Analysis of Variance (ANOVA) tests were used to determine the effect of group (maternal separation, control) on log EcoG power at each EcoG frequency during each of the five behavioral states. Multiple comparisons were adjusted by 5% threshold Benjamini–Hochberg False Discovery Rate (FDR) (Benjamini and Hochberg, 1995). FDR-corrected p-values of less than 0.05 were considered statistically significant. Statistical analyses were performed using the rstatix package in R (version 4.02).

## Results

### Maternal separation paradigm and home cage behavioral assessment used for EcoG analysis and preprocessing

We sought to assess effects of an established early-life stress paradigm, 3-hr daily maternal separation and isolation from littermates (MS), on early brain activity as indexed by EcoG power. The experimental timeline began on P0 with the birth of the litters as determined by video monitoring. The 3-hr long daily MS paradigm began on P2 and lasted until P10. The pups were separated from the dam and isolated from their littermates while the dam was kept in a new cage in a separate room. Following the last separation on P10, pups were implanted with telemetry devices to record frontal EcoG activity. From P11 to P14, daily home cage EcoG recording sessions took place with careful monitoring of dam-pup interactions to determine the effect of Arched Back Nursing, Licking and Grooming, Milk Ejections, In-Nest, and Out-of-Nest on pups’ cortical activity. As presented in the following section, there were no group differences in maternal behavior for any of the dam-pup interactions monitored. Overall, we found that MS pups had lower frontal EcoG power relative to Controls.

### MS leads to lower EcoG power spectra during dam-pup behavioral interactions

We first assessed group differences in EcoG power at each frequency during different dam-pup behavioral interactions: Out of Nest (OUT), In-Nest (IN), Arched Back Nursing (A), Milk Ejection (ME) and Licking and Grooming (LG). We also analyzed the percentage of time spent in each of these behaviors in the same pups. There were no significant differences in the percentage of time spent in each dam-pup behavior interactions between MS and C groups (*p* > 0.05). Therefore, we are confident that the percentage of time in each behavior interactions did not account for differences observed in the EcoG data.

A repeated measures ANOVA revealed significant interactions of Group × Frequency Band during most of the behaviors and for the majority of the days analyzed (Fig. 2). Notably, significantly l ower EcoG power was found i n MS compared to C groups on P13 when the dam was OUT, which was primarily observed in the high frequency bands (27-100 Hz). Similarly, MS pups on P13 had lower EcoG power in the high frequency bands during A (25-100 Hz; *p* < 0.05). Group differences were also found during ME on P11, although for this behavior and postnatal day MS pups had significantly lower EcoG power across all frequencies as compared to C. This effect of MS was not found on P12. On P13, during LG, MS pups had lower EcoG power from 5 to 50 Hz. Thus, overall, pups in the MS group had lower EcoG power that was primarily found in the high frequency band range except for LG, where the group effect on EcoG power was also found in the lower frequency bands.

**Fig. 2.**
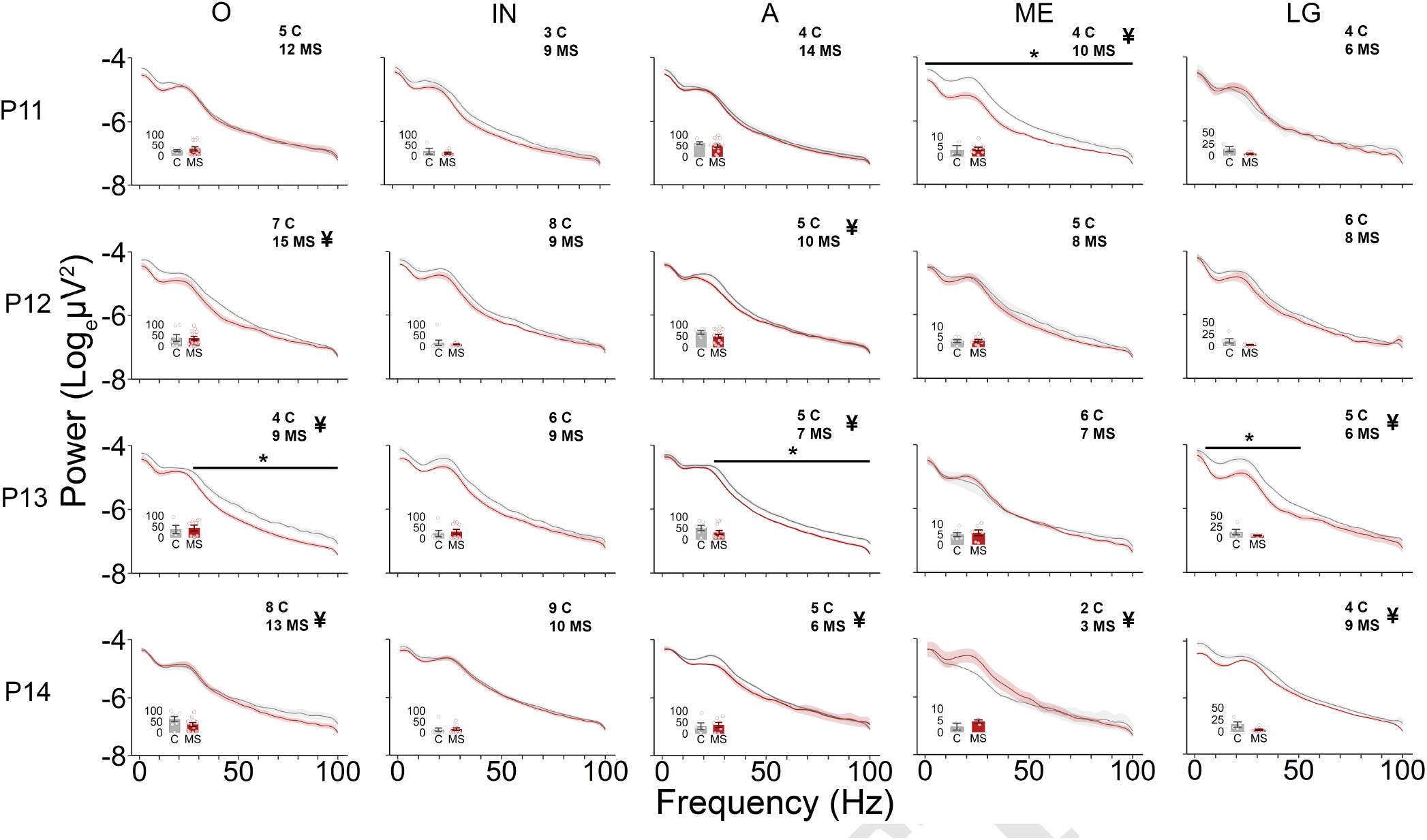
MS leads to changes of EcoG power spectra during important dam-pup behavioral epochs. Average power (log(power)) spectral density curves are shown for each day (P11, P12, P13, P14) for different behaviors (Out of Nest, In-Nest, Arched Back Nursing, Milk Ejection, and Licking and Grooming) of C (grey) and MS (red) rat pups. Out of Nest P11 (Group : *F*_(1,15)_ = 0.113, *p* = 0.74, Band: *F*_(1,99)_ = 308.0, *p* < 0.001, Group × Band : *F*_(1,99)_ = 0.772, *p* = 0.95), Out of Nest P12 (Group : *F*_(1,20)_ = 2.92, *p* = 0.10, Band: *F*_(1,99)_ = 485.1, *p* < 0.001, Group × Band : *F*_(1,99)_ = 1.682, *p* < 0.001), Out of Nest P13 (Group : *F*_(1,11)_ = 9.350, *p* = 0.011, Band: *F*_(1,99)_ = 831.4, *p* < 0.001, Group × Band : *F*_(1,99)_ = 2.282, *p* < 0.001), Out of Nest P14 (Group : *F*_(1,19)_ = 0.940, *p* = 0.35, Band: *F*_(1,99)_ = 252.3, *p* < 0.001, Group × Band : *F*_(1,99)_ = 3.467, *p* < 0.001); In-Nest P11 (Group : *F*_(1,10)_ = 1.847, *p* = 0.20, Band: *F*_(1,99)_ = 484.6,*p* < 0.001, Group × Band : *F*_(1,99)_ = 0.606, *p* = 1), In-Nest P12 (Group : *F*_(1,15)_ = 5.760, *p* < 0.05, Band: *F*_(1,99)_ = 417.9, *p* < 0.001, Group × Band : *F*_(1,99)_ = 0.690, *p* = 0.99), In-Nest P13 (Group : *F*_(1,13)_ = 5.519, *p* < 0.05, Band: *F*_(1,99)_ = 704.3, *p* < 0.001, Group × Band : *F*_(1,99)_ = 0.780, *p* = 0.94), In-Nest P14 (Group : *F*_(1,13)_ = 1.684, *p* = 0.213, Band: *F*_(1,99)_ = 827.6, *p* < 0.001, Group × Band : *F*_(1,99)_ = 0.272, *p* = 1). Arched Back Nursing P11: (Group : *F*_(1,16)_ = 0.517, *p* = 0.48, Band: *F*_(1,99)_ = 668.3, *p* < 0.001, Group × Band : *F*_(1,99)_ = 0.217, *p* = 1), Arched Back Nursing P12: (Group : *F*_(1,13)_ = 0.797, *p* = 0.39, Band: *F*_(1,99)_ = 461.1, *p* < 0.001, Group × Band : *F*_(1,99)_ = 1.142, *p* < 0.01), Arched Back Nursing P13: (Group : *F*_(1,10)_ = 12.35, *p* < 0.01, Band: *F*_(1,99)_ = 1.124, *p* < 0.001, Group × Band : *F*_(1,99)_ = 1.398, *p* < 0.01), Arched Back Nursing P14: (Group : *F*_(1,8)_ = 12.35, *p* < 0.32, Band: *F*_(1,99)_ = 334.8, *p* < 0.001, Group × Band : *F*_(1,99)_ = 1.499, *p* < 0.01); Milk Ejection P11 (Group : *F*_(1,12)_ = 22.23, *p* < 0.001, Band: *F*_(1,99)_ = 347.0, *p* < 0.001, Group × Band : *F*_(1,99)_ = 2.287, *p* < 0.001), Milk Ejection P12 (Group : *F*_(1,11)_ = 1.193, *p* = 0.30, Band: *F*_(1,99)_ = 328.6, *p* < 0.0.001, Group × Band : *F*_(1,99)_ = 0.415, *p* = 1), Milk Ejection P13 (Group : *F*_(1,11)_ = 0.154, *p* = 0.70, Band: *F*_(1,99)_ = 183.5, *p* < 0.001, Group × Band : *F*_(1,99)_ = 0.694, *p* = .99), Milk Ejection P14 (Group : *F*_(1,3)_ = 0.311, *p* = 0.62, Band: *F*_(1,99)_ = 144.5, *p* < 0.001, Group × Band : *F*_(1,99)_ = 2.34, *p* < 0.001); LG P11: (Group : *F*_(1,5)_ = 0.005, *p* < 0.95, Band: *F*_(1,99)_ = 147.8, *p* < 0.001, Group × Band : *F*_(1,99)_ = 1.271, *p* = 0.054), LG P12: (Group : *F*_(1,10)_ = 1.045, *p* = 0.331, Band: *F*_(1,99)_ = 332.1, *p* < 0.001, Group × Band : *F*_(1,99)_ = 0.521, *p* = 1), LG P13: (Group : *F*_(1,9)_ = 8.821, *p* < 0.05, Band: *F*_(1,99)_ = 387.9, *p* < 0.001, Group × Band : *F*_(1,99)_ = 2.538, *p* < 0.001), LG P14: (Group : *F*_(1,11)_ = 6.998, *p* < 0.05, Band: *F*_(1,99)_ = 663, *p* < 0.001, Group × Band : *F*_(1,99)_ = 1.618, *p* < 0.001); Each figure also includes the following: sample size/group; bar graph inset showing the percentage duration of each behavior that occurred per animals used for EcoG analysis. Note that LG behaviors could co-occur during A (low-arched nursing), therefore LG spectral density curves may include some overlap in EcoG data during A, as well as the bar graph insets; ¥displayed next to sample size indicating significant Group X Frequency Band interaction (FDR-corrected *p* < 0.05); horizontal line with * indicating the range where FDR correction showed significance. EcoG, electrocorticography; MS, maternal separation group; C, Control group; P, postnatal day; LG, licking and grooming. EcoG, electrocorticography; MS, maternal separation group; C, Control group; P, postnatal day; LG, licking and grooming.

### EcoG power changes during behavioral transitions are generally blunted by MS

We next investigated how EcoG power changes during dam behavioral transitions. Repeated measures ANOVA revealed a significant Group × Frequency Band interaction on the percent change in EcoG power on P12 during the transition from Pre-ejection (60s preceding ME) to ME (15s during ME); Figure 3A provides an example spectrogram. Notably, the increase in power during ME was substantial on P12, reaching 300% but was less pronounced on P13 (Fig. 3B). A repeated measures ANOVA on the transition from ME (15s during ME) to Post-Ejection (60s duration after ME) revealed a significant effect of Group; MS pups had a significantly lower change in EcoG power but no Group × Frequency Band interaction was found (Fig. 3C).

**Fig. 3.**
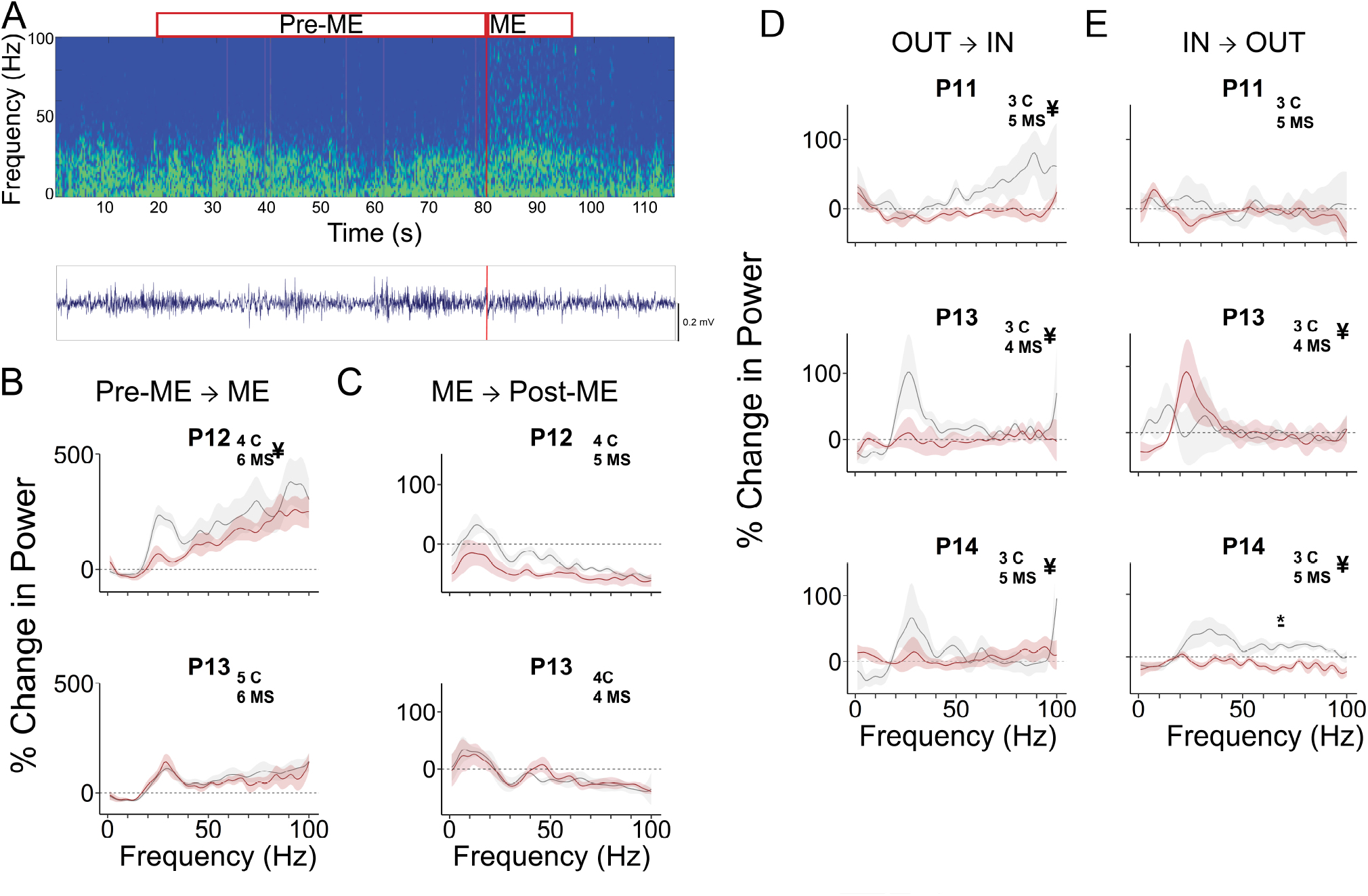
EcoG power changes during behavioral transitions are affected by MS. **(A)** Example sonogram (top) and EcoG trace (bottom) of one rat pup showing the timepoint at which a transition between Pre-Milk Ejection (Pre-ME) and Milk Ejection (ME) took place. Red bound boxes show the duration of the recording which were analyzed. Grey lines show epoch containing artifact that would be removed during pre-processing. **(B)** Average percent change in EcoG power (% Change in Power) during a transition from a 60-s Pre-ME to 15-s ME on P12 (top) and P13 (bottom) across all frequencies. A significant Group × Frequency Band interaction was revealed on P12. High frequency power increased up to 300% in C pups on P12. Pre-ME → ME, P12: Group *F*_(1,5)_ = 0.348, *p* = 0.58, Band *F* (1, 99) = 6.113, *p* < 0.001, Group × Band *F*_(1,99)_ = 1.306, *p* < 0.05. An increase in high frequency EcoG power was also observed on P13. Pre-ME → ME P13: Group *F*_(1,6)_ = 0.185, *p* = 0.68, Band *F*_(1,99)_ = 10.64, *p* < 0.001, Group × Band *F*_(1,99)_ = 0.599, *p* = 0.99). **(C)** Average percent change in EcoG power (% Change in Power) during a transition from a 15-s ME to 60-s Post-Milk Ejection (Post-ME) on P12 (top) and P13 (bottom). Overall significant Group and Frequency Band but no interactive effects were observed. ME → Post-ME P12: Group *F*_(1,7)_ = 6.377, *p* < 0.05, Band *F*_(1,99)_ = 7.129, *p* < 0.001, Group × Band *F*_(1,99)_ = 0.859, *p* = 0.83; ME → Post-ME P13: Group *F*_(1,6)_ = 0.014, *p* = 0.91, Band *F*_(1,99)_ = 7.968, *p* < 0.001, Group × Band *F*_(1,99)_ = 0.435, *p* = 1. **(D)** Average percent change in EcoG power (% Change in Power) during a transition from a 60-s Out of Nest (OUT) to In-Nest (IN) on P11 (top), P13 (middle) and P14 (bottom). OUT → IN: P11 Group *F*_(1,5)_ = 6.723, *p* < 0.05, Band *F*_(1,99)_ = 2.343, *p* < 0.001, Group × Band *F*_(1,99)_ = 1.48, *p* < 0.01; OUT → IN: P13 Group *F*_(1,6)_ = 3.07, *p* = 0.13, Band *F*_(1,99)_ = 2.373, *p* < 0.001, Group × Band *F*_(1,99)_ = 1.703, *p* < 0.001; OUT → IN: P14 Group *F*_(1,9)_ = 0.076, *p* = 0.79, Band *F*_(1,99)_ = 1.657, *p* < 0.001, Group × Band *F*_(1,99)_ = 1.909, *p* < 0.001. **(E)** Average percent change in EcoG power (Change in Power) during a transition from a 60-s IN to OUT on P11 (top), P13 (middle) and P14 (bottom). IN → OUT: P11 Group *F*_(1,6)_ = 0.275, *p* = 0.62, Band *F* (1, 99) = 0.685, *p* = 0.99, Group × Band *F*_(1,99)_ = 0.832, *p* = 0.87); IN → OUT P13 Group *F*_(1,5)_ = 0.005, *p* = 0.95, Band *F*_(1,99)_ = 1.588, *p* < 0.001, Group × Band *F*_(1,99)_ = 1.407, *p* < 0.05; IN → OUT: P14 Group *F*_(1,7)_ = 16.557, *p* < 0.01, Band *F*_(1,99)_ = 3.286, *p* < 0.001, Group × Band *F*_(1,99)_ = 2.11, *p* < 0.001. Lines and shaded confidence intervals are displayed by group, MS in red and C in grey. Dashed line indicates 0% change in EcoG. ¥ displayed next to sample size indicate significant Group × Frequency Band interaction (FDR-corrected *p* < 0.05); horizontal line with ∗ indicate the range where FDR correction showed significance. EcoG, electrocorticography; MS, maternal separation group; C, Control group; P, postnatal day; LG, licking & grooming.

On P11, P13 and P14 when dam was transitioning from Out of Nest (OUT) to In Nest (IN), repeated measures ANOVA revealed a significant Group × Frequency Band interaction effect on EcoG power (*p* < 0.01; Fig. 3D). On P13 and P14, we also found a significant Group × Frequency Band interaction on the reverse transition from In to Out (*p* < 0.05; Fig. 3E). On P14, a significant Group and Group × Frequency Band interaction were observed; post-hoc testing showed a significant difference in the 67-70 Hz frequency range, where MS pups had lower EcoG power compared to Controls (FDR-corrected *p* < 0.05). Together these results suggest that MS had an overall blunting effect on the changes in EcoG power during behavioral transitions, especially for those involving the dam moving in and out of the nest. That is, EcoG power for pups in the MS group showed smaller changes in response to dams’ transitions as compared to controls.

### MS does not affect EcoG activity during the maternal potentiation paradigm

We examined whether the MS rearing condition affects EcoG power during the three phases of the potentiation paradigm (ISO1, REUN, ISO2; Fig. 4). There were no significant effects of Group or Group × Frequency Band interactions on EcoG power during any of the phases of isolation or reunion (Fig. 4B). Furthermore, when comparing the phases to each other in terms of % change in power, there were no significant differences observed irrespective of rearing condition. These results suggest that MS rearing condition may affect pups’ EcoG power only when in the normal home cage situation.

**Fig. 4.**
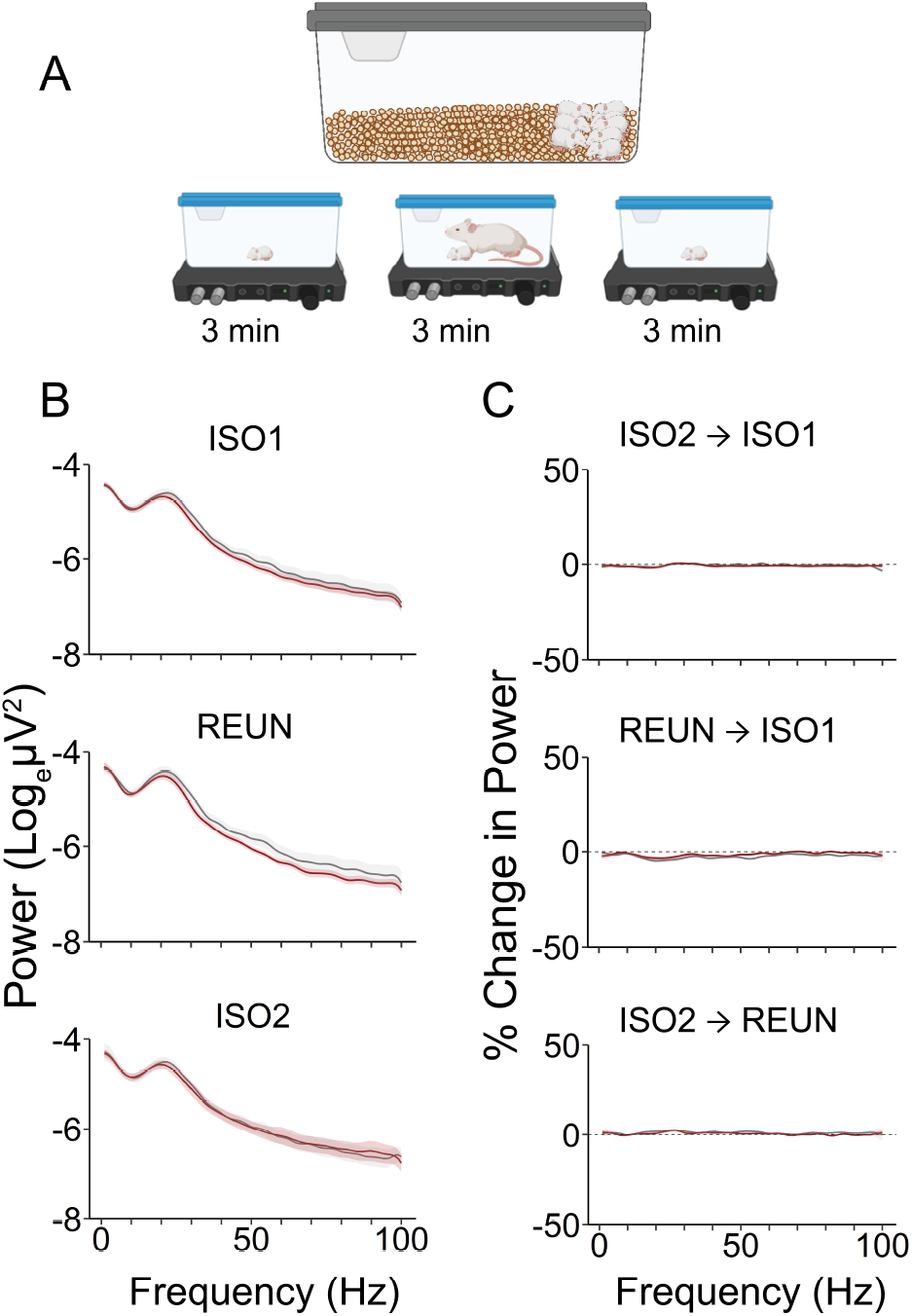
MS does not affect EcoG activity during maternal potentiation paradigm. **(A)** Potentiation paradigm. After a 30-min separation from the dam in the home-cage, each pup is removed and placed alone in an novel small clear rectangular box for 3 minutes (ISO1). The dam is reunited with the pup for 3 minutes (REUN) and then removed from the box for a second 3-min isolation (ISO2). **(B)** Average power spectra during each phase. No significant differences were found in any of our variables of interest. **C** Percent change in EcoG power (% Change in Power) between ISO2 and ISO1 (top), REUN and ISO1, and ISO2 and REUN for Group and Group × Band were not significant, but a significant Band effect was found. ISO2 → ISO1: Group : *F_P_* _(1,18)_ = 0.261, *p* = 0.62, Band *F*_(1,99)_ = 2.564, *p* < 0.001, Group × Band *F*_(1,99)_ = 0.771, *p* = 0.95; REUN → ISO1: Group *F*_(1,20)_ = 2.218, *p* = 0.15, Band *F*_(1,99)_ = 3.45, *p* < 0.001, Group × Band *F*_(1,99)_ = 0.396, *p* = 1; ISO2 → REUN: Group *F*_(1,19)_ = 1.504, *p* = 0.24, Band *F*_(1,99)_ = 1.365, *p* < 0.05, Group × Band *F*_(1,99)_ = 0.298, *p* = 1. EcoG, electrocorticography; MS, maternal separation group; C, Control group; ISO1, first isolation period; REUN, reunion between dam and pup; ISO2, second isolation period.

## Discussion

### Fronto-cortical responses are generally blunted in MS pups during normal behavioral interactions and transitions

Here we present the first demonstration that MS from P2-P10 affects fronto-cortical EcoG power and power changes in the pre-weaning period (P11-P14) during normally occurring dam-pup behavioral interactions and transitions. Observed differences between MS and normally reared pups varied by postnatal day, behavior/behavioral transition and frequency band, but overall MS had a blunting effect on both power and power changes with two exceptions. First, MS pup fronto-cortical activity was higher in mid-frequency bands (15-40 Hz) on P14 during milk ejection. This is particularly interesting in light of the starkly lower power observed across all frequency bands in this group during milk ejection at P11. Second, MS pup fronto-cortical power increased in mid-frequency bands (20-30 Hz) during dam transitions from in-nest to out-of-nest on P13. No power change was observed in normally reared pups during this transition and interestingly, the opposite was observed during dam transitions from out-of-nest to in-nest, where we observed increased power in the mid-frequency band in normally reared pups but not MS pups. Overall, power differences between MS and normally reared pups were larger when the dam was out-of-nest than when the dam was in-nest. In terms of nurturing behaviors, the blunting effect on EcoG power peaked earlier for arched-back nursing and milk ejection (P12-P13 and P11, respectively) than for licking and grooming (P13-P14). Interestingly, these differences in fronto-cortical activity during behavioral transitions did not extend to our dam-pup reunion-isolation paradigm, in which we previously reported MS to significantly impact multiple acoustic features of USVs (Kaidbey et al., 2019).

Prior studies have showed that male MS rats, both during the juvenile period while anesthetized (Reincke and Hanganu-Opatz, 2017) and during slow wave sleep in adulthood (Mrdalj et al., 2013) have significantly lower frontal EEG power compared to normally reared controls. Here, we show the origins of this fronto-cortical change to occur already at P11-P14 during naturally occurring behaviors.

Our observations of fronto-cortical EcoG power in rat pups during normal home cage behavioral interactions align with recent findings from other groups using neocortical LFP recordings (Sarro et al., 2014; Courtiol et al., 2018). We note that while others have opted for classification of brain electrical signals into bands, such as delta, theta, alpha, beta and gamma, here we restricted our analysis to actual frequencies for two reasons: the wide variability in the literature on exact definitions of these bands and the lack of physiological evidence of clear-cut relevance of these bands to neural activity and behavior. Nevertheless, some consensus exists on low versus high frequency bands and their contribution to various states. For example, increased power at high frequencies (gamma, >30 Hz) is associated with active sleep in both animals and humans (Llinas and Ribary, 1993; Joliot et al., 1994; Maloney et al., 1997; Honjoh et al., 2018), while “attentive” waking is associted with higher power in high frequencies (>30 Hz) but also in the theta band (6-10 Hz) (Jones, 2020). Conversely, quiet (non-rapid eye movement [REM]) sleep is associated with increased power at low frequencies (delta, 0.5−4 Hz) (Jones, 2020). Prior work showed rat pup neocortical LFP power in high frequency bands (beta and gamma frequency ranges 24–72 Hz) is increased when the dam is out of the nest (Sarro et al., 2014) while LFP power in low frequency bands (delta/theta: 0−8 Hz) was greater when pups were attached to their dam’s nipple preceding a milk ejection, i.e. during nurturing contact with the dam (Courtiol et al., 2018). It was proposed that maternal caring behavior concurrently and positively modulates cortical brain activity to enhance synchronicity in normally reared pups (Sarro et al., 2014; Courtiol et al., 2018). Here, we replicate this effect during maternal presence and care provided to normally reared rat pups, and extend this finding by showing less pronounced changes in frontal EcoG low frequency power in MS pups, as well as lower EcoG power in high frequency bands in MS pups when the dam is out of the nest.

Our results also align with recent findings using a different rodent model of early life stress. In a model of scarcity-adversity (i.e., twice daily 1 hour bouts of low-bedding between P10-P14), rat pup neocortical LFPs showed similar dampening of power across most frequencies when dam was exhibiting nurturing maternal behaviors (e.g. licking and grooming, milk ejections) compared to normally reared controls (Opendak et al., 2020). Interestingly, in this model, minimal impact was observed on the pup neocortical LFPs during the actual rough handling. This aligns with our own finding that EcoG power differences were most pronounced during similar nurturing behaviors and not observed during overall dam in-nest time. Additionally, the profound differences between stressed and unstressed (control) rat pups in high frequency cortical responses to transitions (e.g., during milk ejection) that we and others have reported are possibly indicative of changes in cortical arousal. Thus, it appears that high stress-inducing rearing conditions (i.e., MS, adversity-scarcity) affect normal cortical maturation manifested by cortical hyporeactivity.

### Possible mechanisms underlying MS-mediated fronto-cortical activity changes

It is not clear how MS dampens cortical reactivity. One hypothesis is that lower EcoG power in MS pups results from neuronal dysmaturation. Juvenile rats with a history of MS have structural and functional neuronal abnormalities in the medial prefrontal cortex (Chocyk et al., 2013). Myelination in the prefrontal cortex goes through an inverted “U”-shape trajectory over the lifespan in rats, with extremely low levels of myelination prior to P7, a slow increase to weaning (P21), and a rapid increase peaking at P90 (Yang et al., 2017). Impairments in myelination in the medial prefrontal cortex of young mice and rats exposed to MS has been previously reported (Yang et al., 2017; Teissier et al., 2019).

Another potential mediator is altered maturation of the gamma-amino-butyric acid (GABA) system. GABA is critical to several developmental processes, including regulation of cell differentiation, cell migration, dendritic arborization, and synapse formation and stabilization (Ben-Ari et al., 2007; Wang and Kriegstein, 2008). Disruptions in GABA function during sensitive periods of development result in structural and functional changes leading to dysfunction in sensory processing, emotion regulation, and cognition (Li et al., 2002). Reduced maternal care and pain exposure has been previously associated with alterations in frontal and hippocampal GABA-levels and GABA-to-glutamate relative levels (i.e. altered excitation/inhibition balance) in 4-day old rat pups (Mooney-Leber et al., 2018). Therefore, EcoG power differences observed in the MS pups in our model could reflect GABA-mediated imbalances in excitatory/inhibitory drive due to delayed (or accelerated) developmental transition from the excitatory to the inhibitory actions of GABA (Ben-Ari et al., 2007).

It is interesting to note that this developmental shift in GABA effects on neurons is thought to be facilitated by oxytocin signaling (Khazipov et al., 2008; Leonzino et al., 2016). Given the role of oxytocin release and regulation by mother/infant interactions (Welch and Ludwig, 2017; Vittner et al., 2018; Markova and Siposova, 2019), repeated separation from the dam might lead to overall lower central oxytocin release, which, in turn, might delay normal maturation of GABA inhibitory circuitry.

### A model for understanding early-life stress effects in infants

The explicit goal of this study was to generate a model that recapitulates our observations in NICU infants with and without FNI, a nurture-based behavioral strategy to enhance maternal-infant emotional connection shown to improve developmental trajectories of these infants (Welch et al., 2012, 2013). The most prominent early findings we reported in FNI infants were increased EEG power during sleep and improved cortical functional connectivity compared to standard care infants (Welch et al., 2014; Myers et al., 2015; Welch et al., 2017). The most significant effects were localized to the left frontal polar region (Welch et al., 2014; Myers et al., 2015). Comparable to standard care infants in the NICU, who receive relatively less maternal care than the FNI group, we here showed that MS pups have blunted left fronto-cortical activity compared to normally reared pups. This suggests that fronto-cortical activity can be used as an early marker, and possible mediator, of long-term effects of this early-life stressor (i.e. chronic maternal separation). Further exploration in this rat paradigm will elucidate mechanisms of maladaptive and adaptive brain responses to MS, and help develop novel interventions for protecting NICU infants, a highly vulnerable population.

## ACKNOWLEDGEMENTS

This work was sponsored by generous donations from Fleur Fairman, John and Rainy Erwin, and Einhorn Collaborative.

## Supplementary Information

**Fig. 5.**
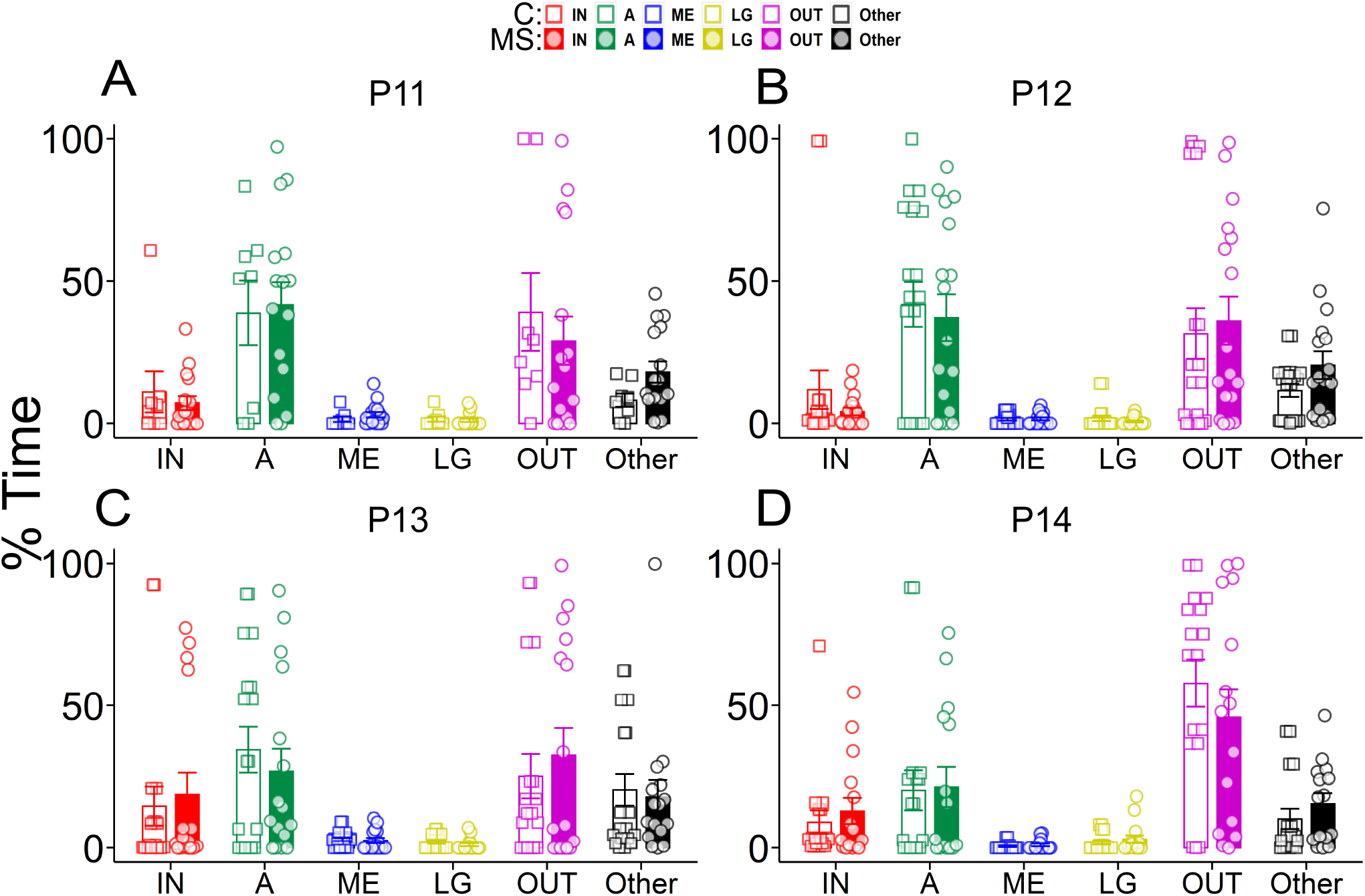
Bar graph showing the behavioral epoch durations expressed as a percentage of total recording time (% Time) for each daily recording, from P11 to P14, with standard error bars and distribution of each data points. Sample sizes for each recording day are as follows: P11: C (*n* = 12), MS (*n* = 16); P12: C (*n* = 20), MS (*n* = 17); P13: C (*n* = 18), MS (*n* = 16); P14: C (*n* = 17), MS (*n* = 15). MS, maternal separation group; C, Control group; P, postnatal day.

